# Workflow for multiplex microsatellite panel development and sample preparation for robust amplicon sequencing of low-template and degraded DNA: validation for non-invasive genotyping in three large carnivore species

**DOI:** 10.64898/2026.08.20.745956

**Authors:** Marta De Barba, Frédéric Boyer, Molly Baur, Marjeta Konec, Elena Pazhenkova, Nadège Remollino, Céline Stoffel, Barbara Boljte, Christian Miquel, Tomaž Skrbinšek, Pierre Taberlet, Luca Fumagalli

## Abstract

High-throughput amplicon sequencing has transformed microsatellite (STR) genotyping by overcoming many of the limitations of fragment-length analysis, enabling more accurate, cost-effective, and standardized genotyping. Yet, protocols specifically designed for high-throughput sequencing (HTS)-based STR genotyping from low-template and degraded DNA remain scarce, despite the prevalence of these challenging sample types in ecological and conservation contexts. We present a methodology for the *de novo* development of robust STR multiplex panels together with a laboratory protocol for efficient and reliable STR genotyping by sequencing with low quantity and quality DNA samples. The protocol comprises (i) an automated bioinformatic pipeline to design large sets of short tetranucleotide markers optimized for multiplex amplicon sequencing of degraded and low-template DNA; (ii) guidelines for efficient *in vitro* optimization of multiplex amplification using directly low quantity/quality template DNA; and (iii) a library preparation procedure that improves detection of low-level allele signal while enabling quality assessment of STR amplicon sequencing under limiting DNA conditions. We demonstrate the approach by developing and validating STR panels for non-invasive genotyping of three large carnivore species: a 44-plex for the grey wolf (*Canis lupus*), a 41-plex for the Eurasian lynx (*Lynx lynx*), and a 30-plex for the brown bear (*Ursus arctos*). Multiplex performance was high, with ≥91% of samples successfully genotyped at ≥50% of loci (allele size range 28–110 bp across panels) and correctly assigned to known individuals, negligeable levels of noise in the controls, and high discriminatory power (PIDsibs ≤ 2.4 × 10^−12^), also owing to sequence variation among same-length alleles at 15–50% of loci. The approach is broadly applicable to animal and plant species, a wide range of sample types, and large-scale analysis such as genetic monitoring. Our study reinforces the value of STR amplicon sequencing for ecological and conservation applications while highlighting the importance of marker design and laboratory workflows tailored to HTS-based genotyping for accurate and efficient implementation.

## Introduction

The advent of high-throughput sequencing (HTS) technologies (Reuter, Spacek & Snyder, 2015) has revolutionized the field of molecular ecological research by advancing genomic approaches and facilitating marker development also for non-model species (Ekblom & Galindo, 2011; Porter & Hajibabaei, 2018), therefore widening the application and impact of molecular tools for biodiversity conservation (Hohenlohe, Funk & Rajora, 2021; Theissinger et al., 2023; Schiebelhut et al., 2024). One significant outcome of the increasingly widespread use of HTS has been the improvement of genotyping of commonly used markers in population genetics and genetic monitoring, such as microsatellites, also known as short tandem repeats (STRs), and single nucleotide polymorphisms (SNPs), through genotyping-by-sequencing approaches (Campbell, Harmon & Narum, 2015; Vartia et al., 2016; Baetscher et al., 2018). These methods employ HTS-based amplicon sequencing of markers designed from genome-wide data for enabling high multiplexing levels and offer advantages over approaches based on higher marker density and locus coverage, as they allow time- and cost-effective genotyping of large sample sizes and technical ease of implementation in standard genetic laboratories (Meek & Larson, 2019; Krehenwinkel et al., 2019).

HTS of STRs was developed in response to the need of overcoming limitations of classical STR genotyping based on laborious fragment analysis on capillary electrophoresis platforms. The main advantage of STR amplicon sequencing consists in providing direct access to the STR sequences which allows for (i) accurate scoring of STR alleles based on both nucleotide sequence and length polymorphism, including the detection of new allelic variants or mutations; (ii) standardization and automation of the genotyping process through bioinformatics; and (iii) data comparability across laboratories and platforms (Fordyce et al., 2015; De Barba et al., 2017; Curto et al., 2019). These features, together with high data throughput, higher multiplexing capacity, increased sensitivity to low DNA quantities and decreased cost of massive sequencing, have made the method highly suitable for large sample sizes (Krehenwinkel et al., 2019). Since the early studies showing successful implementation and the potential for ecological applications (Suez et al., 2016; Vartia et al., 2016; Darby et al., 2016; Zhan et al., 2017; De Barba et al., 2017), workflows and analytical tools for STR genotyping by HTS have been further developed. Procedures for STR multiplex panel development and HTS implementation (Curto et al., 2019; Krehenwinkel et al., 2019; Lepais et al., 2020), as well as software and bioinformatic pipelines for automated genotyping (Hoogenboom et al., 2017; Zhan et al., 2017; Barbian et al., 2018; Roy et al., 2021; Huo et al., 2021; Wang et al., 2023; De Barba et al., 2024; Liu et al., 2024; Perry et al., 2024) are now available and have promoted the adoption of this approach across diverse ecological systems. In less than 10 years, HTS of STR markers has been applied across plants, vertebrates, and invertebrates to investigate phylogeographic patterns, genetic diversity and structure, gene flow, and population abundance, as well as for genetic assignment, parentage, and mixture analyses (Bradbury et al., 2018; Blondel et al., 2019; Curto et al., 2019; Feyrer et al., 2019; Ruzzante et al., 2019, 2020; Krehenwinkel et al., 2019; D’Aloia et al., 2020; Li et al., 2021; Lepais et al., 2022; Allegue et al., 2024; Jiang et al., 2024; Beimfohr et al., 2025).

STRs are particularly valuable and have been among the most widely used markers for genotyping low-quantity and highly degraded DNA in non-invasive genetic studies and monitoring, as well as in analyses of forensic and archival plant and animal specimens (Andrews et al., 2018; Carroll et al., 2018; Nakahama, 2021; Sahajpal, Mishra & Bhandari, 2021; Ramírez-García et al., 2025; Sandikci, 2026). Although SNP genotyping has also been optimized for low quantity/quality DNA using microfluidic (Kraus et al., 2015; von Thaden et al., 2017; Nygaard et al., 2022) and more recently GT-seq (Eriksson, Ruprecht & Levi, 2020; Burgess, Irvine & Russello, 2022; Hayward et al., 2022; Hervey et al., 2025) approaches, STRs remain the preferred marker system for a broad range of applications requiring genetic profiling from challenging samples. In addition to robust amplification under limiting DNA conditions, the high allelic diversity of these markers confers increased information content per locus, enabling improved resolution even in populations with limited genetic diversity and for the detection and interpretation of mixed DNA samples containing genetic material from multiple individuals. By comparison, other genetic and genomic approaches require substantially more loci and correspondingly larger amounts of input DNA to achieve comparable levels of statistical power (Andrews et al., 2018; Eriksson, Ruprecht & Levi, 2020; Gang & Shrivastav, 2021). Further, their high polymorphism and relatively low ascertainment bias allow use and reliable diagnostic testing across a species’ full range (Harpending & Rogers, 2000). HTS of multiallelic STRs, based on the determination of allele sequences, can significantly enhance these applications by providing increased genotyping accuracy, success, efficiency and standardization (De Barba et al., 2017). However, most studies using STR amplicon sequencing for genotyping poor DNA quantity/quality sources have so far focused on testing and validation (Donaldson et al., 2020; Trede et al., 2021; Salado et al., 2021; Walker et al., 2026), with only few population-level studies (Yuan et al., 2022; Wolfenson et al., 2024) and no routine large-scale applications published so far.

A critical factor for successful and efficient implementation of HTS of STRs from low quality/quantity DNA samples is the selection of optimal markers, which requires more stringent criteria compared with conventional genetic samples containing high DNA quality and quantity. Marker features that should be prioritized with these sources are (i) short length of amplified products (<150 bp) for ensuring robust amplification of degraded DNA; (ii) ability to co-amplify with a limited amount of DNA template, to enable high level of PCR multiplexing in laboratory processing; (iii) reduced strand slippage during PCR to minimize PCR-induced stutter sequences (often more pronounced with challenging DNA sources) and enable accurate and automated allele calling using bioinformatics (De Barba et al., 2017).

This latter characteristic is particularly important for amplicon sequencing, as loci producing high levels of artifactual sequences may result in inaccurate genotyping and need significant manual verification, which is also the main reason for favoring tetra-nucleotide versus shorter repeat motifs. In addition, existing STR assays are suboptimal for HTS-based genotyping due to larger amplicon size heterogeneity and a higher tendency for missing data and scoring errors, compared to STRs developed *de novo* for amplicon sequencing and designed *ad hoc* for high multiplexing (Lepais et al., 2020). Consistent with this, studies evaluating HTS-based genotyping of existing markers used for capillary electrophoresis genotyping, including short repeat motifs such as di-nucleotides, have reported high levels of sequencing artifacts and error rates for genotyping DNA from fecal and museum samples (Donaldson et al., 2020; Yuan, Malekos & Hawkins, 2021; Salado et al., 2021). While various protocols have been proposed for marker development (e.g. Curto et al., 2019; Krehenwinkel et al., 2019; Guo et al., 2020; Lepais et al., 2020; Antunes et al., 2022), none specifically address the full range of constraints associated with low-template and degraded DNA sources.

In addition to marker development and selection, bench protocols are of paramount importance when working with low-quantity and degraded DNA, as they significantly affect sequencing performance and genotyping success. Particularly, the adoption of appropriate PCR products purification and library preparation methods can minimize the occurrence of spurious sequence reads and the noise-to-allele ratio in HTS outputs from these DNA sources, which typically show higher background noise than good quality DNA samples. As a result, these approaches considerably increase the number of viable reads for genotyping and improve allele detection sensitivity and scoring accuracy under limiting DNA conditions. For example, multiple cleanup steps can be important to remove non-targeted sequences resulting from amplification of degraded DNA from noninvasive samples, as well as free residual adapters or index-primers in multiplexed sequencing libraries that could cause index misassignment and result in false inferences (Eriksson, Ruprecht & Levi, 2020). Similarly, PCR-free library preparation protocols have been shown to be very effective for reducing the occurrence of tag-jumps causing incorrect assignment of sequence reads to samples in DNA metabarcoding data from mixtures of nucleotide-tagged amplicons (Carøe & Bohmann, 2020) and individual genotyping of environmental DNA (eDNA) samples (De Barba et al., 2024).

Adoption of standardized laboratory procedures tailored to maximize accurate detection of low-level allele signal from low-quantity and degraded DNA samples, as well as the use of appropriate internal controls to evaluate the performance of STR amplicon sequencing, are therefore essential for quality assurance and efficient implementation of HTS STR genotyping in large-scale ecological and forensic studies.

In this study, we present a validated methodology for developing robust multiplexing panels and a bench protocol for efficient and reliable sequencing-based STR genotyping of samples with low DNA quantity and quality. The approach is designed for broad applicability across animal and plant species and is scalable to large-size projects. We applied the methodology to *de novo* marker development and genotyping in three large carnivore species, the grey wolf (*Canis lupus*), the Eurasian lynx (*Lynx lynx*) and the brown bear (*Ursus arctos*), using samples collected in the field such as scats, saliva from prey remains, and urine to demonstrate its effectiveness for ecological applications involving degraded and low-quantity DNA.

### Material & Methods

### STR marker development and multiplex optimization

Here we present the protocol in a generalized form to facilitate broader applicability. In a section below, we describe its adaptation for application to three large carnivore species. The development of the genotyping assay is composed of four steps (Fig. 1).

1. The initial step consists of identifying from shotgun or genome assembly data multiple STR loci suitable for HTS-based genotyping of low-quantity and degraded DNA (De Barba et al., 2017), that is (i) tetranucleotide loci due to lower stutter artifact rates facilitating automated analysis; (ii) markers resulting in a total amplicon length (including primers) below 150 bp to maximize amplification efficiency of fragmented DNA and allow complete coverage of the STR array on both strands with short reads (i.e. 2×150 bp) HTS platforms; (iii) primers enabling high PCR multiplexing to increase efficiency of laboratory processing. To meet these conditions, the shotgun or genome assembly sequence data is searched for perfect tetranucleotide STRs with 9-16 repeats (to minimize the likelihood of loci with low polymorphism and arrays exceeding 150 bp) not containing GG or CC in the motif, and for which PCR primers can be identified with the following characteristics: 3’-end positioned as close as possible (max distance 12 bp) to the repeat array, 17-25 bp length, 57±1 °C salt adjusted melting temperature (Tm), not ending with either aa/at/ta/tt, not containing a fragment of ≥10 bp with only 2 nucleotides (e.g. atatttatat) or of ≥5 bp with only cg (e.g. cgggc), not showing low complexity (i.e. all 10 bp subsequences of the primers must have a Shannon entropy of 2 bp motifs >1.1), not containing the locus motif or any di-, tri-, tetranucleotide motif repeated three times. These steps were implemented for automation in a python script available at https://gricad-gitlab.univ-grenoble-alpes.fr/leca/PRIMA. From this initial set of STR loci, a subset of 200 markers is randomly selected for *in vitro* testing.
2. The 200 markers selected from step (1) are tested for co-amplification in four distinct multiplex PCRs of 50 markers each (specifically, multiplex 1 including markers 1-50; multiplex 2 markers 51-100; etc.), using a commercial multiplex PCR kit and equal primer concentrations for all markers. A single DNA extract, or a small pool of DNA extracts from low-quantity/quality biological samples, is used for the selection of markers that provide robust amplification under limiting DNA conditions. Amplifications are carried out taking into account the Tm of the primers and sufficient PCR cycles to amplify low-quality and low-quantity DNA. All amplification products from the four multiplexes are then combined in the same library to be sequenced on a HTS platform. The sequencing output is analyzed to select 50 markers that amplify low quantity/quality DNA in a single multiplex PCR with the highest amplification performance, i.e. resulting in high number of reads of the target sequence and low non-specific amplifications. The number of 50 STRs can be adapted but based on our experience it provides an appropriate starting number of markers for optimizing large multiplexes (i.e. 30-40 co-amplified markers at the end of the process).
3. The STR markers selected in the previous step are then combined to evaluate their amplification performance in a single multiplex PCR. At this stage, primer concentrations in the PCR are readjusted, based on the number of reads obtained in the previous step, i.e. higher concentrations for markers yielding fewer reads and lower concentrations for markers yielding more reads. The same DNA samples initially tested are used, but separate PCR amplifications and sequencing libraries are prepared for each sample. PCR conditions and removal of poorly performing markers are as described in step (2). This step may be repeated, with adjustments to the primer concentrations of the markers retained, until the number of marker reads is appropriate and balanced across markers; PCR replicates may also be performed to assess repeatability.
4. This step corresponds to the final marker validation for genetic tagging applications in ecological and population genetic studies and forensics. It involves amplicon sequencing of an adequate number of field samples with PCR replication, and allele calling using available bioinformatic pipelines to infer sample genotypes (see references in the Introduction). Amplification success, genotyping success and error rates, polymorphism and power for individual identification are assessed, and tests for Hardy-Weinberg equilibrium, linkage disequilibrium and null alleles performed. This step may lead to the exclusion of additional few markers failing to meet validation criteria and entail further tuning of primer concentrations to accommodate for the final set of retained markers and their performance across larger sample sizes with variable DNA quality/quantity.

**Fig. 1:**
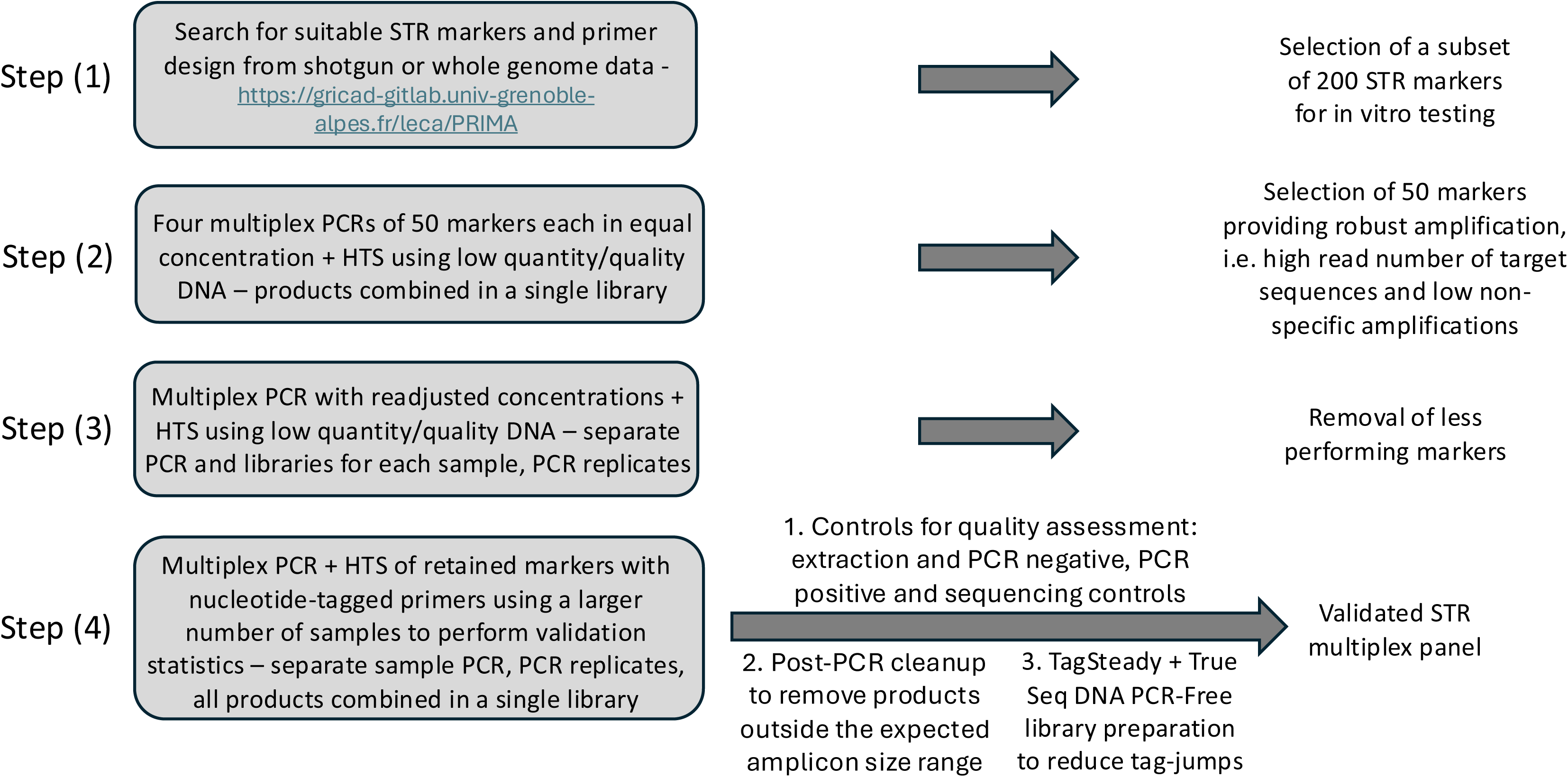
Diagram of the workflow describing the STR development steps, genotyping protocol and outcome.

### HTS-based STR genotyping protocol

We describe a HTS genotyping protocol based on 150 bp paired-end (PE) sequencing of pooled nucleotide-tagged amplicons using Illumina or Element Biosciences platforms, which is particularly convenient for routine analysis of large sample sizes in ecological projects (Taberlet et al., 2018). This general workflow can be adapted to other sequencing protocols and platforms.

#### Multiplex PCR set up using nucleotide-tagged primers

The optimal working unit for STR genotyping based on amplicon sequencing using nucleotide-tagged primers is a 96-well PCR plate to process 96 samples simultaneously, including biological samples and controls. In this setup, a single PCR plate corresponds to one PCR replicate for each sample in the plate. The multitube approach (Taberlet et al., 1996) is applied for ensuring reliable genotyping of poor quality/quantity DNA by performing several independent PCRs for all samples, e.g. eight 96-well PCR plate replicates with the full multitube approach. PCR primers in the optimized multiplex are modified by the 5’-addition of molecular identifier tags, allowing the assignment of sequences to samples and markers in post-sequencing bioinformatic analysis. Tags consist of eight nucleotides enabling a minimum of five mismatches between any pair of tags (Taberlet et al., 2018). For example, with 32 forward and 24 reverse tags, a total of 32 x 24 = 768 unique tag combinations are possible on eight PCR plates (Appendix), a strategy suitable for large-scale assays (Taberlet et al., 2018). An additional 1–3 specified nucleotides are added to the tags 5’-end to increase complexity for enhanced cluster detection on the sequencing flow cell. Using this approach, multiplexed products from the eight PCR plates can be efficiently sequenced together as a single library pool.

To facilitate set up of the genotyping multiplex PCR using nucleotide-tagged primers with large sample sizes, we recommend preparing three distinct plate types: *DNA aliquot plate*, *Master mix plates*, and *Primer plates*.

The *DNA aliquot plate* is a 96-well plate containing aliquots of the *n* DNA samples intended for PCR amplification. In addition to the biological samples, a *DNA aliquot plate* must also include the extraction negatives processed during DNA extraction of the biological samples (unless verified before), and a number of PCR negative, PCR positive, and tagging system controls (Fig.1, Appendix). These controls are necessary as the resulting sequence data will be used to detect possible contamination at different stages of the process, monitor the performance of the amplification and the sequencing, and set filtering parameters in the bioinformatic pipeline (De Barba et al., 2014, 2024). PCR negatives can be made by sterile water (at least 2 recommended per plate), while positive controls are DNA samples whose genotype has previously been determined (at least 2 recommended per plate). We stress the importance of using positive control samples with a quality index (Miquel et al., 2006) resembling an average field sample, to balance the number of resulting reads. Tagging system controls or “blanks” correspond to empty wells on the PCR plate (i.e. no PCR mix, primers, and template), allowing estimation and monitoring of tag jumps (Schnell, Bohmann & Gilbert, 2015) which can significantly decrease the number of viable reads available for genotyping (8-12 recommended per plate, Appendix). Blanks should be disposed diagonally on the plate to allow checking for the highest possible number of unused tagged combinations, ideally one blank per forward and reverse tag (Taberlet et al., 2018, Appendix).

*Master mix plates* are 96-well PCR plates containing all PCR reagents except primers and DNA. A master mix for *n* amplification reactions can be prepared using a commercial multiplex PCR kit following manufacturer’s instructions and then distributed across the *Master mix plates* corresponding to independent PCR replicates based on the *DNA aliquot plate* design. *Primer plates* are 96-well PCR plates containing all possible combinations of forward and reverse tagged primers of the multiplexed markers. *Primer plates* (1X concentrated) are prepared by first pooling forward and reverse primers carrying the same tag into separate 2X primer mixes (one mix for each forward tag containing all forward primers and one mix for each reverse tag containing all reverse primers). These mixes are then distributed into 96-well plates such that each well contains a unique combination of forward- and reverse-tagged primers for all markers in the multiplex (see the Appendix for illustration).

To set up the genotyping multiplex PCR, aliquots from the *DNA aliquot plate* and the P*rimer plates* are added to the *Master Mix plates* according to the final DNA and primer concentrations in the PCR. Thermocycling is carried out following the multiplex PCR kit manufacturer’s instructions and talking into account the melting temperature of the markers.

#### Preparation of the pooled library and sequencing

PCR products from all plates are pooled together and purified using the MinElute PCR purification kit (Qiagen GmbH) for post-PCR cleanup, allowing elution in a suitable volume for library preparation. A second purification is subsequently conducted using SPRI beads (Beckman Coulter) to remove unspecific PCR products exceeding the amplicon size threshold established in step (1) (i.e. 150 bp) (Fig. 1). Finally, the concentration of the purified pool is measured (Qubit 4 Fluorometer, Thermo Fisher Scientific), and a fragment analysis (Advanced Analytical) is performed to evaluate the size and relative abundance of the amplicons.

To minimize tag jumps, i.e. low-level detection of alleles from different samples because of false DNA tag combinations that appear as experimental artifacts, it is essential to prepare the sequencing library using a PCR-free library preparation method. To this end, we combined the TagSteady protocol (Carøe & Bohmann, 2020), a PCR-free library preparation procedure for pools of nucleotide-tagged amplicons that enables generation of data with virtually no tag-jumps, with the Illumina True Seq DNA PCR-Free library preparation kit (Fig. 1). The Reaction Booster mix and the End-repair master mix are prepared following Carøe & Bohmann (2020). A 10 µl aliquot of the latter is added to 2 pmol of purified PCR products resuspended in 20 µl, resulting in a final reaction of 30 µl. This mixture is then incubated in a thermocycler, according to the conditions specified in the TagSteady protocol. From this point onward, the procedure transitions to the True Seq DNA PCR-Free library preparation kit (Illumina), beginning with the “Ligate Adapters” step. A post-library fragment analysis and qPCR (KAPA Library Quantification Kit-Illumina, Roche) are then performed to assess library size and concentration before sequencing.

### Application to three species of large carnivores

To illustrate the methodological approach, we applied the protocol for STR marker development and HTS genotyping to non-invasively collected samples from three large carnivore species: grey wolf (*Canis lupus*), Eurasian lynx (*Lynx lynx*) and brown bear (*Ursus arctos*).

#### Samples and DNA extraction

Samples used in this study included tissue samples obtained from a legally removed wolf in Switzerland and a lynx that died in a vehicle collision in Slovenia, together with non-invasive genetic samples collected in the field during genetic monitoring. Non-invasive samples included wolf scats, saliva, and urine from the Alpine population in Switzerland, lynx scats from the endangered Dinaric population in Slovenia and Croatia, recently reinforced through translocation of individuals from the Carpathian population (Krofel et al., 2025), and brown bear scats from the Dinaric–Pindos population in Slovenia (Table S1).

Tissue samples were collected and stored in 95% ethanol. Scats were sampled by taking ∼1 cm^3^ of scat material and stored either in 95% ethanol or DET buffer (Frantzen et al., 1998). Saliva samples were collected on wolf preys using swabs and stored dry. Urine samples were collected in snow and stored according to Valière & Taberlet (2000). No ethical approval was required to work with non-invasively collected samples or tissues from dead animals (naturally or legally deceased animals). Fieldwork procedures were specifically approved by relevant regional and national wildlife offices as a part of national monitoring activities.

Additional details on the wolf, lynx and brown bear populations and genetic monitoring are available in Dufresnes et al., (2019), Pazhenkova et al., (2025), and Skrbinšek et al., (2019) respectively.

Genomic DNA was extracted from ∼25 mg of tissue material and the saliva swab using the DNeasy Blood and Tissue Kit (Qiagen GmbH), and from 0.1–0.2 mL of feces using the Qiamp Fast DNA stool Mini kit (Qiagen GmbH) or the MagMax DNA Multi-Sample Kit (Thermo Fisher Scientific) whole blood protocol optimized to be used on a liquid handling robot Hamilton Starlet. Urine DNA extraction was carried out according to Valière & Taberlet (2000). A negative control was included in each extraction batch to monitor contamination. DNA extractions of non-invasive samples were performed in a room dedicated to low quantity/quality DNA sources.

#### STR development and validation for grey wolf, Eurasian lynx, and brown bear

For the wolf and the lynx, we used the DNA extracted from the tissue samples to perform shotgun sequencing. The shotgun library was prepared with a TruSeq DNA PCR-Free Library Prep Kit (Illumina), and DNA sequencing was performed using the Illumina MiniSeq platform with the High Output Kit yielding 25 million 150 bp PE reads. For the brown bear, we obtained published whole genome shotgun data from SRA (PE reads for samples SAMEA4762870, SAMEA4762871, SAMEA4762872 of study ERP109572, Barlow et al., 2018).

For each species, genomic data were used for STR identification as described in step (1) of the marker development workflow described above. Step (2) was then accomplished using a mixture of DNA extracted from five scat samples. We used the Platinum Multiplex PCR kit (Thermo Fisher Scientific) to amplify 200 randomly selected markers in four distinct multiplex PCRs of 50 markers each. Amplifications were performed in a total volume of 20 µl, including 2 µl of template DNA, 1 x Platinum Multiplex PCR Master Mix, 0.16 mg/ml of bovine serum albumin (BSA, Roche Diagnostics), and 0.02 µM of each forward and reverse primer. Thermocycling conditions were as follows: denaturation at 95°C for 2 min, followed by 45 cycles of 30 s at 95°C, 30 s at 55°C (2°C less than the melting temperature), 1 min at 72°C, with a final elongation step of 7 min at 72°C. The pool of all multiplex products for each species was sequenced on a single Illumina library. Library preparation was conducted with a TruSeq DNA PCR-Free Library Prep Kit (Illumina), and DNA was sequenced on the Illumina MiniSeq platform, using the Mid Output Kit generating 8 million 150 bp PE reads, with 15% PhiX. We used the OBITools (Boyer et al., 2016) and simple Unix commands to sort the sequence output by marker and select 50 best performing markers for each species.

The selected markers were co-amplified in a single multiplex PCR (step 3), readjusting primer concentrations within a range of 0.025 µM to 0.035 µM and performing separate amplifications and sequencing libraries for each sample. After evaluating marker performance, we repeated this step using the retained markers, carrying out four replicates per sample and readjusting primer concentrations within a less stringent range of 0.02 µM to 0.045 µM. While doing this, we ensured that the sum of the concentrations of all forward and reverse primers in the multiplex fell within 2-4 µM, as recommended by the manufacturer (Thermo Fisher Scientific). One library was prepared and sequenced for each sample replicate. Sequence outputs were analyzed as above.

Marker validation (step 4) was done by using the final set of STRs and optimized primer concentrations for each species to genotype 76 wolf (33 scats, 40 saliva, 3 urine), 42 lynx (scats), and 86 brown bear (scats) samples (Table S1). We used the HTS genotyping protocol described above with nucleotide-tagged primers, except performing multiplex amplifications for the lynx and the bear in a volume of 10uL with 1uL of template. Samples were selected to represent DNA of varying quality based on previous individual genotyping and included biological replicates of the same animal as an additional assessment of genotyping repeatability. Wolf and lynx samples were sequenced on an Element Biosciences Aviti platform (70 and 44 million PE reads, respectively), whereas brown bear samples were sequenced on an Illumina MiniSeq (14 million PE reads).

The sequence data output of each species was processed to infer STR alleles from the sequences and their relative read counts in each PCR product using the process and automated pipeline (https://github.com/PazhenkovaEA/ngs_pipelines.py/) described in De Barba et al., (2017, 2024). Genotypes were then organized in a custom Microsoft Access database that was used to determine consensus genotypes at each locus, perform matching of sample genotypes, and assign sample genotypes to individuals (Skrbinšek et al., 2019, Appendix). Finally, for each species STR panel, we assessed (i) locus amplification success, as the average proportion of locus PCR yielding reads assigned to at least one allele; (ii) locus genotyping success, i.e. the proportion of consensus genotypes obtained for each locus; (iii) matching probabilities (Waits, Luikart & Taberlet, 2001); (iv) number of locus differences between individual multilocus genotypes; (v) per locus genotyping error rates for allelic dropout (ADO) and false alleles (FA) estimated on individually genotyped samples (Broquet & Petit, 2004). Based on the individual genotypes determined, we then examined marker polymorphism, tested for Hardy-Weinberg (HWE) and linkage disequilibrium (LD) and estimated null allele frequencies. All calculations were done in Genalex (Peakall & Smouse, 2006) and in R (R Core Team, 2023) using packages adegenet (v 2.1.11; Jombart, 2008), pegas (v1.4; Paradis, 2010), poppr (v 2.9.8; Kamvar, Tabima & Grünwald, 2014) and PopGenReport (v 3.1.3; Adamack & Gruber, 2014) as described in the Appendix.

## Results

### New STRs for non-invasive HTS genotyping of grey wolf, Eurasian lynx, and brown bear

Shotgun sequencing of the tissue samples generated a total of 31,228,218 2×150 bp reads for the wolf, and of 33,228,592 2×150 bp reads for the lynx. From these and the published brown bear genomic data, we identified 5,715 (*C. lupus*), 1,775 (*L. lynx*) and 2,143 (*U. arctos*) primer pairs flanking a perfect tetranucleotide repeat, meeting the prerequisite selection criteria (step 1). Testing a subset of 200 markers resulted in 0-1,048,001 (wolf), 0-444,768 (lynx), and 0-1,002,353 (bear) reads/marker, from which 50 markers (read count ≥2,165) were selected for each species (step 2). These were combined for co-amplification into a single multiplex, resulting in the selection of 44 (wolf), 41 (lynx), and 30 (bear) best performing markers to be included in the final multiplex (Table 1), with an average number of reads/marker/PCR across samples of 6,772 for wolf, 30,132 for lynx, and 24,289 for bear (Step 3).

**Table 1:**
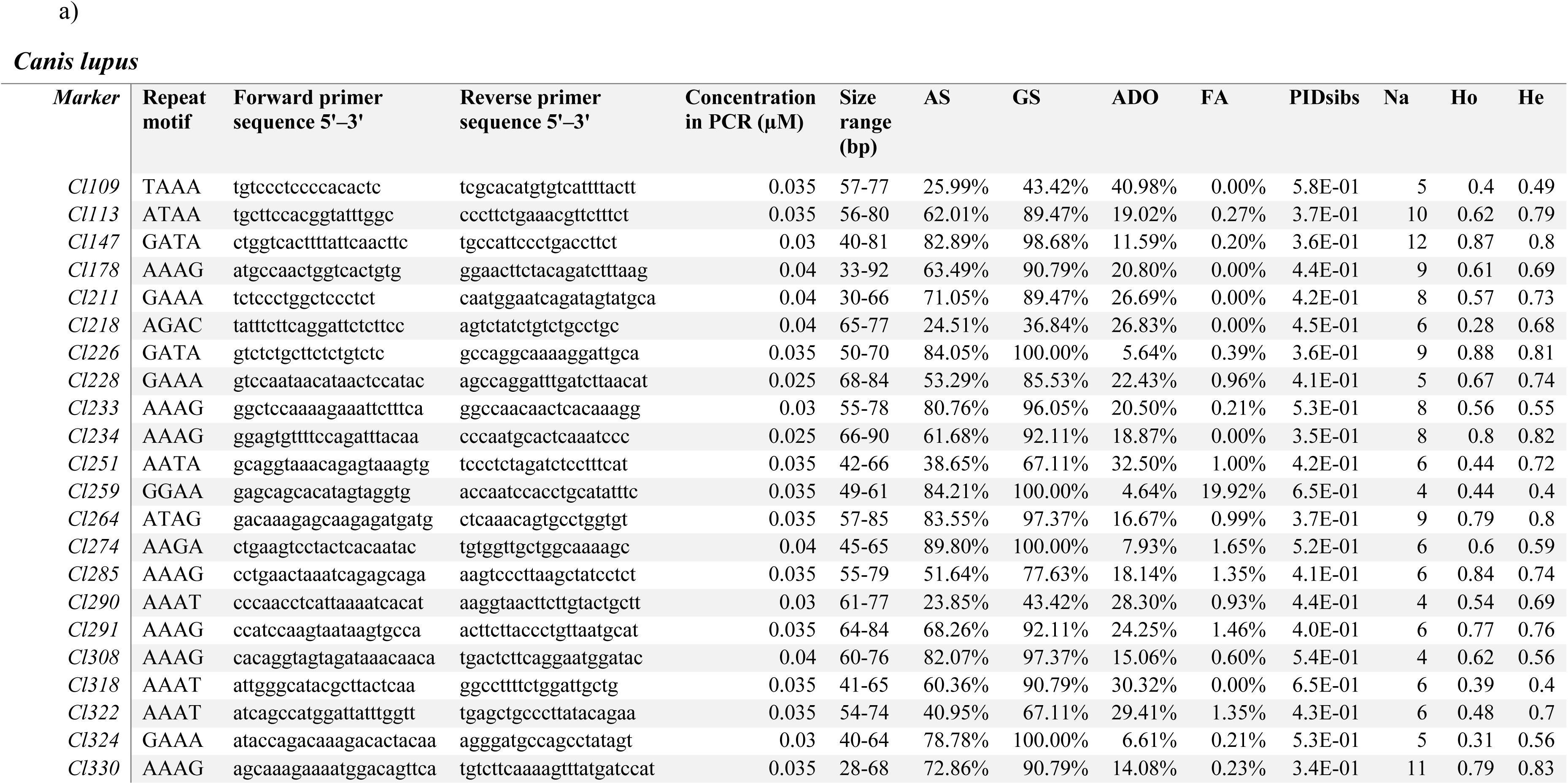
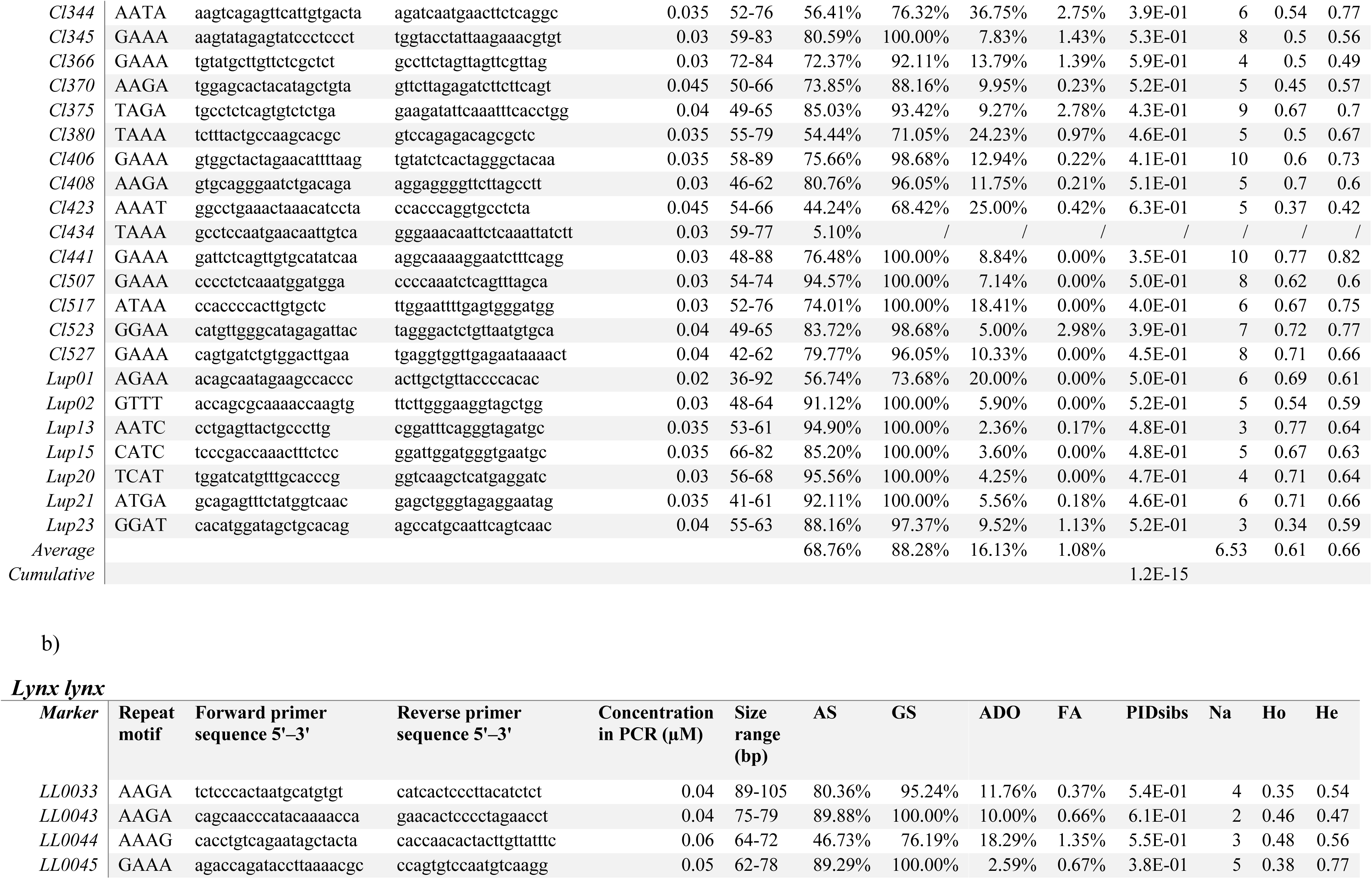
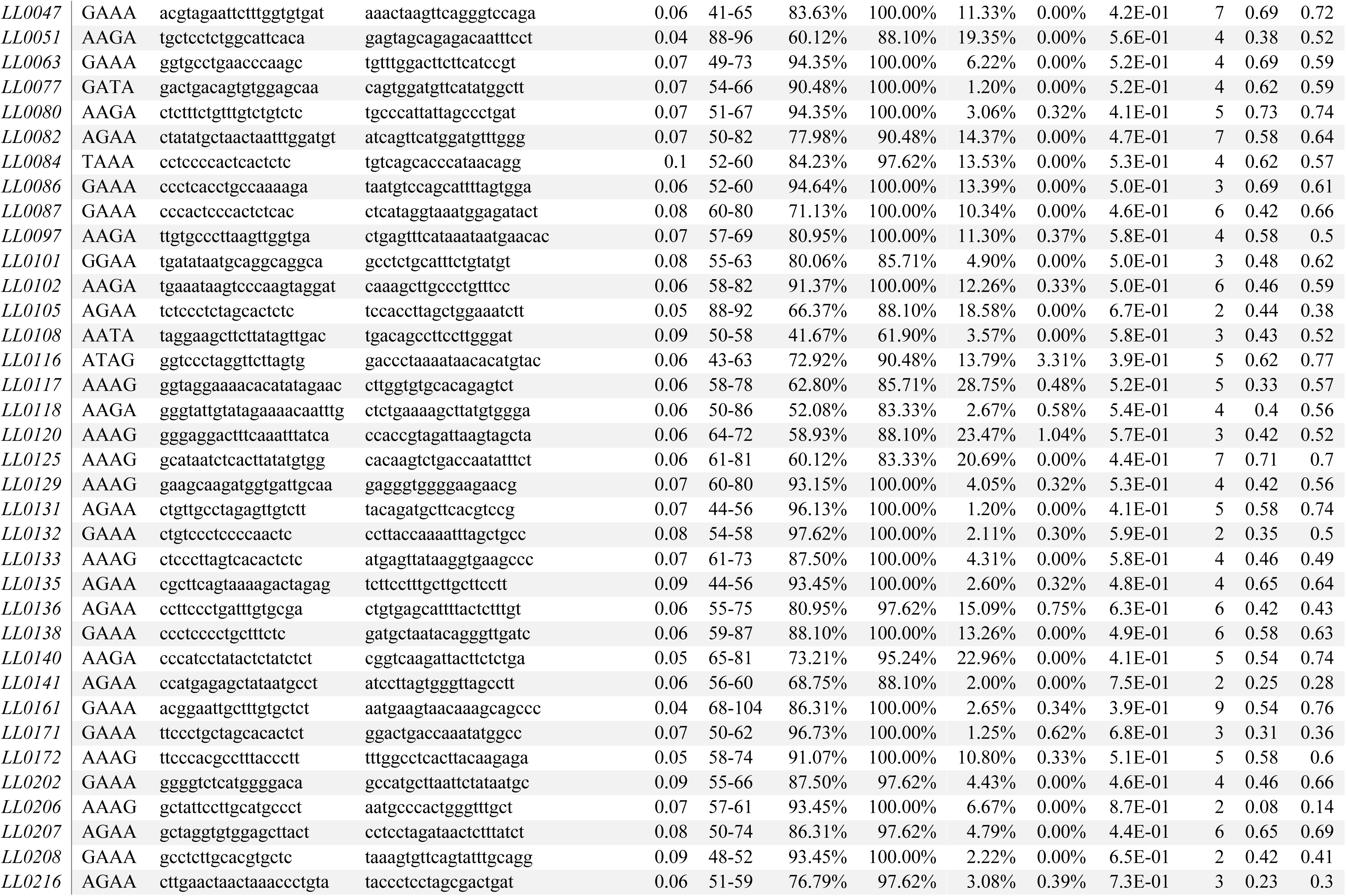
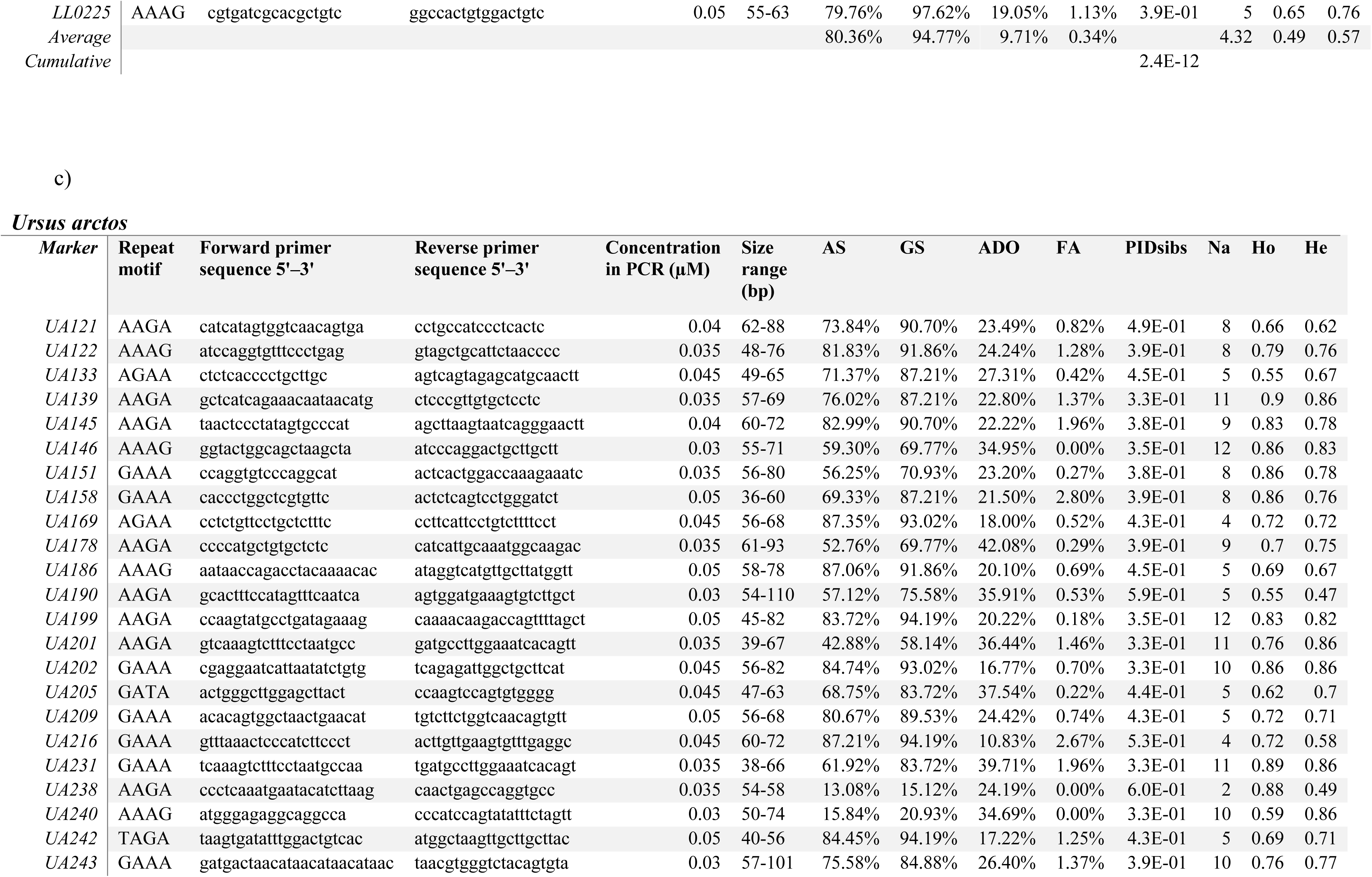
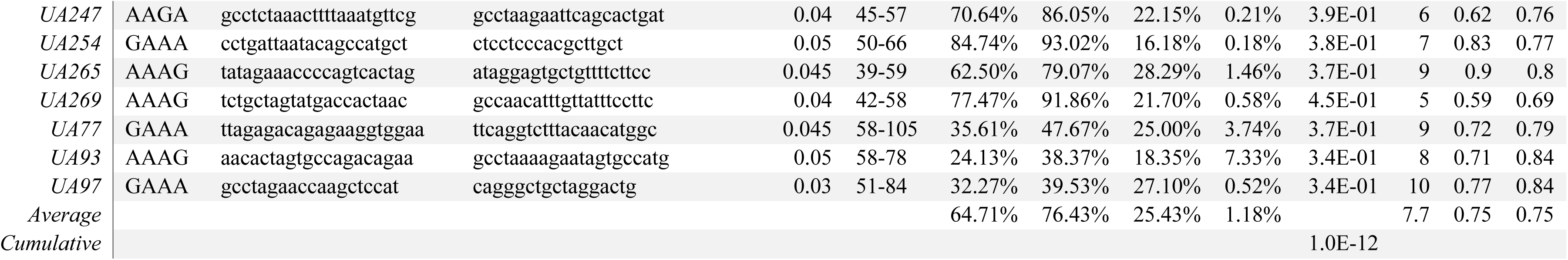
Newly developed STR markers included in the final wolf 44-plex (a), lynx 41-plex (b), and brown bear 30-plex (c). A.S: average amplification success; GS: average genotyping success; ADO: allelic dropout; FA: false alleles; PIDsibs: probability of identity among siblings; Na: number of alleles; Ho: observed heterozygosity; He: expected heterozygosity.

### Marker panel validation

The newly developed multiplex panels were used for genotyping non-invasive samples for the three species according to the multiplex HTS genotyping protocol described above to perform marker validation (step 4). Amplicon sequencing generated 48,207,130 reads for wolf, 11,451,395 reads for lynx, and 7,941,265 reads for brown bear that were assigned to markers and samples, with an average of 11,426 (wolf), 9,859 (lynx), 2,607 (bear) reads/marker/PCR that were used for genotyping. The level of reads observed in the negative and tagging system controls was negligible (0.00003-0.0007%).

The STR loci had an average amplification success of 68.8% in wolves, 80% in lynx, and 65% in bears (Table 1). In wolves, one locus (Cl434) showed only 5% amplification success during validation and was excluded from subsequent analyses. Genotyping success ranged 37–100% (mean 88%) for 43 wolf loci, 62–100% (mean 95%) for 41 lynx loci, and 15–94% (mean 76%) for 30 bear loci (Table 1). All wolf and lynx samples, and 91% of bear samples, were successfully genotyped at ≥50% of loci (22, 21, and 15 loci, respectively; Table S2-4) and were correctly assigned to their known individuals. Across all species, individual genotypes differed at ≥13 loci, cumulative PIDsibs values were ≤ 2.4 × 10^−12^, and average ADO and FA rates ranged 9.71-25.43% and 0.34-1.18% respectively (Table 1).

Genetic diversity was highest in the bear (mean 7.7 alleles per locus, Ho = 0.75, He = 0.75), intermediate in the wolf (6.5 alleles, Ho = 0.61, He = 0.66), and lowest in the lynx (4.3 alleles, Ho = 0.49, He = 0.57) datasets (Table 1). The total observed allele size range for all species markers was 28-110 bp. Size homoplasy, due to SNPs or indels in the repeat unit or the flanking regions, was detected in all three panels, affecting 57 of 281 alleles (20.3%) in wolves, 18 of 177 (10.2%) in lynx, and 66 of 231 (28.6%) in bears, distributed across 17, 6, and 15 loci, respectively (Table S5). Five wolf loci and zero to five lynx loci (depending on whether translocated lynx and their offsprings were included in the analysis) showed significant deviations from HWE, whereas no loci deviated in bears after Holm-Bonferroni correction (Holm, 1979) (P < 0.05; Table S6). No departures from linkage equilibrium were detected in any species after correction (Table S7). Estimated null allele frequencies were below 0.2 for all bear loci, for all wolf loci except four, and for all lynx loci except five (Table S6).

Detailed analysis results of the genotyping validation run for each species are presented in the Appendix.

## Discussion

Ecological and conservation applications often rely on genotyping of DNA sources consisting of degraded and low-template animal and plant DNA, such as non-invasively collected wildlife material, museum and historical specimens, eDNA samples, and forensic traces. Efficient and reliable genotyping of these sample types is mined by lower amplification success, higher levels of missing data, genotyping errors, and an increased risk of contamination compared to standard genetic samples. Consequently, their analysis typically requires methods and protocol optimization, extensive replication, stringent quality control, and careful analysis, resulting in labor-intensive, time-consuming, and often costly workflows (Waits & Paetkau, 2005; Wandeler, Hoeck & Keller, 2007; Alaeddini, Walsh & Abbas, 2010; Taberlet et al., 2018).

We present here a protocol for STR development and sample processing tailored to accurate and efficient high-throughput genotyping of low-quantity and degraded DNA. The general workflow aligns to other published *de novo* STR development strategies for non-model species from HTS data by including bioinformatic mining of genomic sequences to identify candidate co-amplifying STRs and subsequent *in vitro* multiplex optimization but differs in several key aspects due to constraints specific to challenging DNA sources and HTS-based genotyping. The protocol is highly flexible and scalable, making it readily adaptable across animal and plant systems and compatible with a wide range of standard sample types.

Marker development follows principles applied in De Barba et al., (2017) for designing the first set of brown bear STRs for HTS genotyping and is efficiently implemented here through a pipeline that automates and upscales STR discovery and primer selection from genomic data. The pipeline targets short markers of similar length and designs primers with compatible properties. These are critical features to maximize amplification of degraded DNA, which is often fragmented, while reducing marker amplification bias and increasing multiplexing capacity with low template input. In addition, the pipeline targets tetranucleotide loci because they present reduced strand slippage during PCR compared to shorter repeat motifs (Ghebranious et al., 2003), which can result in pronounced PCR-induced stutter sequences especially in low quality samples, preventing allele calling or inflating error rates. This consideration is extremely important for large HTS datasets of amplified STRs, as typically generated in large-scale ecological projects, where enabling accurate and automated allele calling through bioinformatic processing is essential (Fordyce et al., 2015; De Barba et al., 2017; Donaldson et al., 2020). While di-nucleotide STR markers have been used for HTS genotyping (Donaldson et al., 2020; Salado et al., 2021; Yuan et al., 2022; Walker et al., 2026), we recommend that ecological studies relying on large numbers of poor-quality samples prioritize optimally co-amplifying tetranucleotides to maximize genotyping success and data accuracy, while also ensuring efficient data processing.

Marker selection and optimization consist of a dual step multiplexing process involving sequential removal of markers and concentration adjustments based solely on marker amplification performance, evaluated through two main parameters, namely total read counts and relative proportion of artifactual reads corresponding to non-specific amplifications. Polymorphism is not explicitly evaluated at this stage because selection of polymorphic markers is accounted for in the development step by selecting STRs with a minimum of 9 repeats, which is indicative of likely variation in repeat number. This allowed us to carry out the testing on a small number of samples pulled into a single DNA mixture in the first step, followed by separate amplifications, including PCR replication, and sequencing in the second step, thereby greatly reducing initial laboratory processing and costs. In addition, we tested the protocol directly on biological samples of low DNA quantity/quality. As locus amplification performance varies with differing DNA states, this ensures that a high proportion of the selected markers will perform reliably under the range of DNA conditions encountered in field samples. This simple procedure guarantees the efficient selection of a marker set exhibiting the most robust amplification within a single multiplex PCR and the highest specificity (i.e. producing only the target sequence), both of which directly influence the accuracy of allele calling from HTS data (De Barba et al., 2017). The protocol can be easily scaled to test a variable number of markers from those identified by the pipeline, depending on the aimed size of the final multiplex/es. The selected panel is then validated for routine application on a larger number of samples where fine-tuning adjustments to the multiplex conditions can still be made.

In high-throughput amplicon sequencing applications, the sequence data generated can include a variable amount of noise originating from PCR and sequencing errors, other artifacts, and contamination (Callahan et al., 2016; Davis et al., 2018). The impact of such sources of noise is exacerbated when DNA is degraded and present at low concentrations, where stochastic PCR effects, amplification bias, and contamination become increasingly influential (Pompanon et al., 2005). If the noise-to-true sequence ratio in the sequencing output is high, sensitivity for low-level signals can be reduced and accurate sequence detection compromised. In genotyping studies, this may lead to overall lower amplification success and inflated allelic dropout and false alleles rates, due to failure to detect low-read alleles and an increase in artifactual reads, respectively, therefore negatively impacting genotyping success. Thus, it is advisable to prevent as much as possible the occurrence of spurious reads during laboratory processing prior to sequencing. The protocol of sample preparation we described is adapted for this purpose. Specifically, the double amplicon purification procedure removes unspecific PCR products that are too short or exceed the expected amplicon size, and the optimized library preparation combines the specificity of the TagSteady protocol (Carøe & Bohmann, 2020) with the convenience of the Illumina TruSeq DNA PCR-Free procedure. The extremely low level of reads observed in the negative and tagging system controls included in the analysis of the wolf, lynx and bear non-invasive genetic samples supports the efficacy of the employed protocols to minimize overall noisy background. In this perspective, we emphasize the importance of including internal controls, such as extraction negative, PCR negative, PCR positive, and sequencing controls during the experimental setup, allowing evaluation of STR amplicon sequencing performance, identification of anomalies arising at different stages of the process (e.g. contamination, amplification failure, and high levels of tag-jumps), and the definition of appropriate filtering parameters during bioinformatic analysis. High levels of artifacts observed in the controls should be taken as a warning sign of poor experiment performance and warrant caution in data analysis. Such controls are now a standard in other HTS-based amplicon sequencing applications (De Barba et al., 2014; Zinger et al., 2019). Genotyping-by-sequencing studies should also adopt their systematic use and reporting as a quality assurance measure.

We used the protocol to develop and optimize markers for efficient non-invasive genotyping-by-sequencing of three large carnivore species, specifically a 44-plex for wolf, a 41-plex for lynx, and a 30-plex for brown bear. Validation on previously genotyped samples spanning varying DNA quality and quantity confirmed elevated multiplexing performance, with genotyping success consistent with previous analyses and error rates comparable to those reported for non-invasive genetic studies of these species in the same areas using much smaller multiplexes and capillary electrophoresis-based genotyping (De Barba & Waits, 2010; Sindičić et al., 2013; Dufresnes et al., 2019; Skrbinšek et al., 2019), supporting its suitability for population genetic studies. A few loci performed sub-optimally and/or deviated from equilibrium during validation. Such performance patterns are expected with the highly variable DNA conditions of non-invasive field-collected sources, while some departures from equilibrium may be associated to population status and history (for example, lynx samples originate from a genetically depauperated population that has been the subject of population reinforcement; Pazhenkova et al., 2025) and can also occur by chance due to limited sample sizes. Less performing loci may be further optimized by tweaking primer concentrations or excluding them from the multiplex panel upon feedback from broader application. The lower amplification and genotyping success observed with brown bear samples compared to the other species may reflect the overall lower sample quality (Table S1) as well as sequencing depth in the validation run for bears. Increasing sequencing depth will likely reduce error rates and increase genotyping success, especially for marginal samples (De Barba et al., 2024). Nonetheless, each marker set provided high discrimination power and redundant information for reliable individual identification also in presence of missing locus data and possible residual errors in the consensus genotypes, and even with low levels of genetic diversity, as observed in the lynx data. This is particularly relevant for applications entailing processing of large number of non-invasively collected genetic samples from wildlife populations, because sufficient power for various downstream individual-based analyses (e.g. mark-recapture, genetic diversity, genetic structure and gene flow, pedigree reconstruction and relatedness) can be achieved with a single amplicon sequencing run. We also note that the high information content of large multiplexes is often not required for routine analysis, i.e. solely aimed at individual identification, for which a reduced panel would suffice. This could be advantageous with low quantity/quality samples because smaller multiplexes will likely exhibit even higher performance with such DNA sources. For example, after initial validation, we have successfully used a subset of the markers for individual genotyping of wolf and lynx snow track eDNA (De Barba et al., 2024) and for non-invasive genetic monitoring of wolf in Slovenia (Bartol et al., 2023, 2025a) and Switzerland. Similarly, since 2015, we have been using a small 13-plex STR panel previously developed following the same principles (De Barba et al., 2017) for genetic monitoring of brown bears in Slovenia (Skrbinšek et al., 2017; Bartol et al., 2025b). In addition to STRs, and following initial optimization, PCR primers targeting other classes of genetic markers can also be incorporated into the multiplex assay, including mitochondrial DNA markers for species and lineage identification, sex-identification markers, and diagnostic SNP loci associated with specific genetic variants or mutations.

Across all species, we found a substantial amount of size homoplasy (20%, 10%, 29% of wolf, lynx, and bear alleles, respectively) at multiple loci (39% wolf, 15% lynx, 50% bear), i.e. alleles that would appear identical under size-based capillary electrophoresis but differ at the sequence level due to distinct evolutionary trajectories (Putman & Carbone, 2014). Other STR amplicon sequencing studies showed that STR size homoplasy is indeed very common with varying levels in different species, suggesting underestimation of true STR polymorphism using standard fragment analysis based on capillary electrophoresis (Barbian et al., 2018; Šarhanová et al., 2018; Goncalves et al., 2025; Walker et al., 2026). Accessing the underlying STR sequence variation will allow for more accurate estimation of levels of genetic diversity and enhance the resolution of population genetic analysis in natural populations. This is also relevant for conservation genetic studies and monitoring of genetically depauperated populations, which are often constrained by limited discriminatory power and availability of small amounts of degraded DNA. For example, in the data for the endangered Dinaric lynx, 9 alleles would have been masked using capillary electrophoresis analysis of the same STR markers.

Indeed, the generation of sequence-level genotype data represents a major advancement in STR genotyping enabled by genotyping-by-sequencing approaches. Beyond improvements in automation, standardization, and accuracy, access to STR sequence data renders genotypes independent of the DNA sequencing technology used to generate them, including future technologies. This overcomes longstanding challenges in data comparability and transferability across laboratories and analytical platforms, which have historically imposed major limitations on data sharing, integration, reproducibility, and reusability. In turn, this creates new opportunities for transboundary and longitudinal studies, including standardized genetic surveys and population- or metapopulation-level monitoring of highly mobile species, as well as the development of shared genetic databases spanning species’ geographic ranges. Such advances will support timely research and more effective conservation and management initiatives at biologically relevant spatial and temporal scales.

## Conclusions

In this study we described a methodology for the *de novo* development of robust STR multiplex panels together with a laboratory protocol for efficient and reliable STR genotyping by sequencing with low quantity and quality DNA and demonstrated the approach for non-invasive genotyping of three large carnivore species. Although advances in genomics continue to provide unprecedented insights into evolutionary processes, there is growing recognition that analysis of a relatively small number of traditional genetic markers, such as STRs, still remains a widely accessible, cost-effective essential tool for population studies and for informing conservation (Vieira et al., 2016; Fox et al., 2019; Krehenwinkel et al., 2019; Timm, 2020; Hauser, Athrey & Leberg, 2021). In addition, genomic methods typically require large amounts of high-quality DNA, therefore genetic methods based on reduced panels of informative markers are often an obligated choice when working with limiting DNA conditions (Andrews et al., 2018).

HTS of amplified STRs has emerged as a substantially improved alternative to classical STR genotyping and is being increasingly adopted across a wide range of biological systems (see studies cited in the Introduction), with its use expected to expand further in the future. Nevertheless, current trends in STR genotyping are often overlooked, and several studies continue to emphasize the limitations of traditional fragment analysis methods, particularly when comparing STR and SNP genotyping. However, as the HTS-based genotyping approach solves the majority of problems that were plaguing STR genotyping in the past, we argue that the marker and method choice should be pragmatically driven by the research questions and tailored to the specific application and study system while also accounting for available resources (Bertola et al., 2024).

To use this powerful approach to STR genotyping in practice, we need efficient, streamlined development of optimized markers and laboratory protocols specific to HTS-based genotyping. This is what the marker development strategy and bench protocol presented here provide - they establish a rapid, cost-effective development of highly optimized and informative marker panels and a practical and scalable framework for HTS-based STR genotyping for any species. We believe this will facilitate broader adoption of the approach, with real benefits for large-scale ecological and conservation research.

## Supplemental Information

Supplemental Tables: Sample information and genotyping results

Appendix: list of forward and reverse primer tags, *DNA Aliquot Plate* design, preparation of *Primer Plates*, data analysis and results of species marker validation.

## Additional Information and Declarations

### Author contributions

M.D.B. Conceptualization; Validation; Formal analysis; Writing - Original Draft; Visualization; Supervision; Funding acquisition;

F.B. Software; Formal analysis; Data Curation; Writing - Review & Editing

M.B. Validation; Investigation; Writing - Review & Editing

M.K. Validation; Formal analysis; Investigation; Writing - Review & Editing

E.P Software; Formal analysis; Data Curation; Writing - Review & Editing

N.R. Validation; Investigation; Writing - Review & Editing

C.S. Validation; Investigation; Writing - Review & Editing

B.B. Investigation; Writing - Review & Editing

C.M. Investigation; Writing - Review & Editing

T.S. Validation; Formal analysis; Investigation; Writing - Review & Editing; Funding acquisition;

P.T. Conceptualization; Methodology; Formal analysis; Writing - Review & Editing;

L.F. Conceptualization; Methodology; Validation; Formal analysis; Writing - Original Draft; Visualization; Supervision; Funding acquisition;

### Data availability

Sample list and metadata, forward and reverse primer tags, allele sequences and marker testing results: available in the online Supplemental Information.

Python script for marker development: https://gricad-gitlab.univ-grenoble-alpes.fr/leca/PRIMA.

Raw sequence reads are deposited at the European Nucleotide Archive (ENA study PRJEB122678).

Python and R scripts used for bioinformatic analysis of sequence data together with ngsfilter files for demultiplexing are available from https://doi.org/10.5281/zenodo.21623376.

## Acknowledgements

Some of the lynx noninvasive samples used for protocol validation were collected by the Croatian LIFE LYNX project team.

## APPENDIX List of 32 forward tags and 24 reverse tags

The actual tags are indicated with capitalized letters, and the additional nucleotides to be added are indicated with low cases. The same collection of tags must be added to all primers of the multiplexed markers.

| FORWARD TAGS | REVERSE TAGS |
| --- | --- |
| **1** tACACACAC | **1** ccaACACACAC |
| **2** agACAGCACA | **2** aACAGCACA |
| **3** aacGTGTACAT | **3** atGTGTACAT |
| **4** aTATGTCAG | **4** acaTATGTCAG |
| **5** aaTAGTCGCA | **5** aTAGTCGCA |
| **6** aagTACTATAC | **6** agTACTATAC |
| **7** aACTAGATC | **7** ataACTAGATC |
| **8** aaGATCGCGA | **8** aGATCGCGA |
| **9** aacCGCTCTCG | **9** atCGCTCTCG |
| **10** aGTCGTAGA | **10** atgGTCGTAGA |
| **11** agGTCACGTC | **11** cGTCACGTC |
| **12** acaGACTGATG | **12** agGACTGATG |
| **13** cAGACTATG | **13** acaAGACTATG |
| **14** aaGCGTCAGC | **14** aGCGTCAGC |
| **15** aatTGACATCA | **15** aaTGACATCA |
| **16** cACATGTGT | **16** tgcACATGTGT |
| **17** acGTACGACT | **17** cGTACGACT |
| **18** accATGATCGC | **18** cgATGATCGC |
| **19** cACGACGAG | **19** cgaACGACGAG |
| **20** agCATCAGTC | **20** cCATCAGTC |
| **21** acgATCAGTCA | **21** ctATCAGTCA |
| **22** aTCTACTGA | **22** actTCTACTGA |
| **23** atGATGATCT | **23** tGATGATCT |
| **24** tgtCTGCGTAC | **24** ctCTGCGTAC |
| **25** tAGCGACTA |  |
| **26** acTCAGTGTC |  |
| **27** agaACTCTGCT |  |
| **28** cATATAGCG |  |
| **29** cgCTATGCTA |  |
| **30** accTCGCGCTG |  |
| **31** cAGCACAGT |  |
| **32** atTAGCTAGT |  |

### *DNA aliquot plate* design

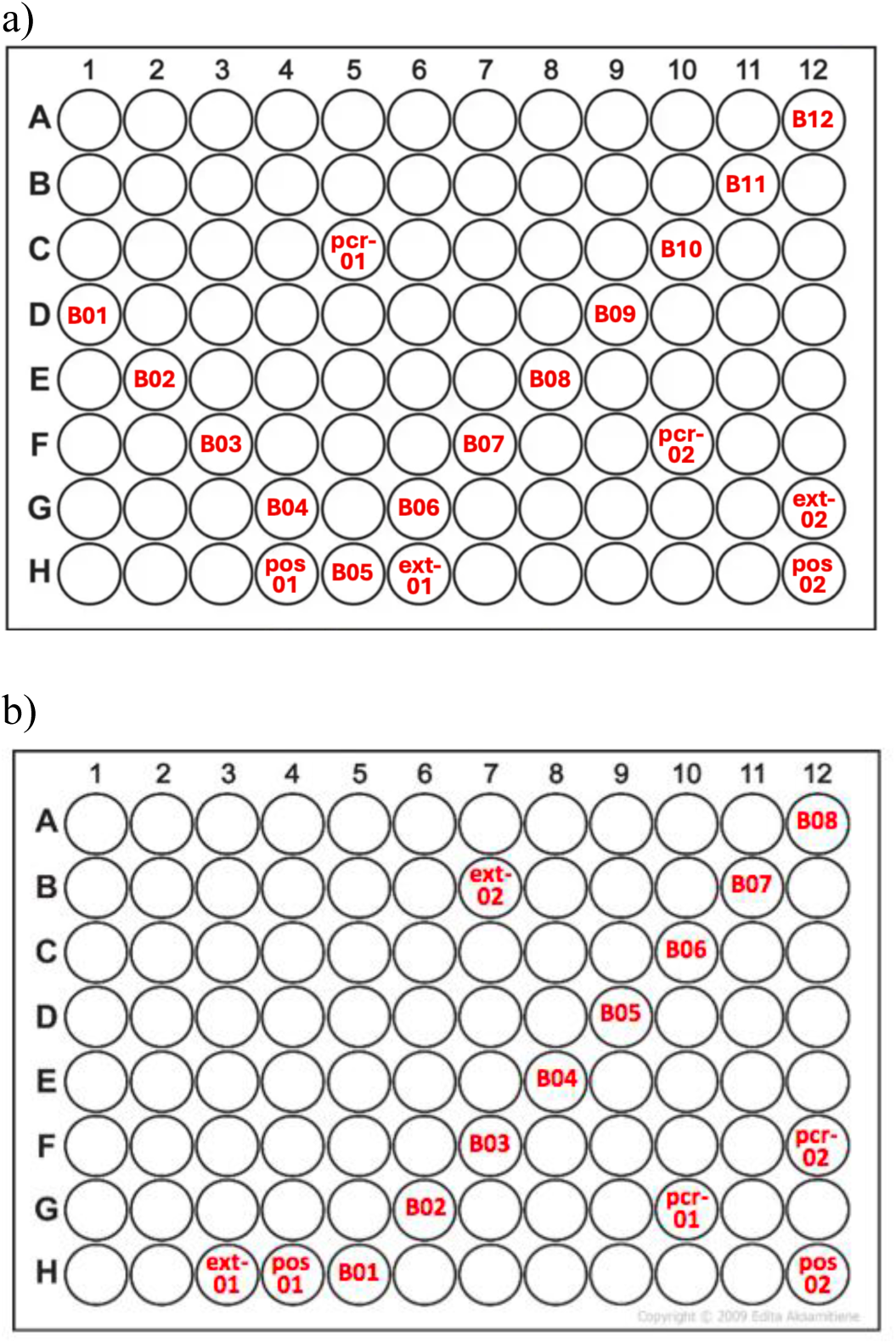

Examples of *DNA aliquot plate* design. B = tagging system controls (blanks), ext- = extraction controls, pcr- = PCR controls, pos = positive controls. a) Full blank design: 12 blanks disposed diagonally on the plate to check all possible unused tagged combinations, i.e. one blank per forward and reverse tag; b) Reduced blank design: 8 blanks disposed diagonally on the plate allowing to check a subset of unused tagged combinations, i.e. some forward or reverse tags are not covered by the blanks, allowing to include a higher number of biological samples in the plate.

### Preparation of *Primer plates*

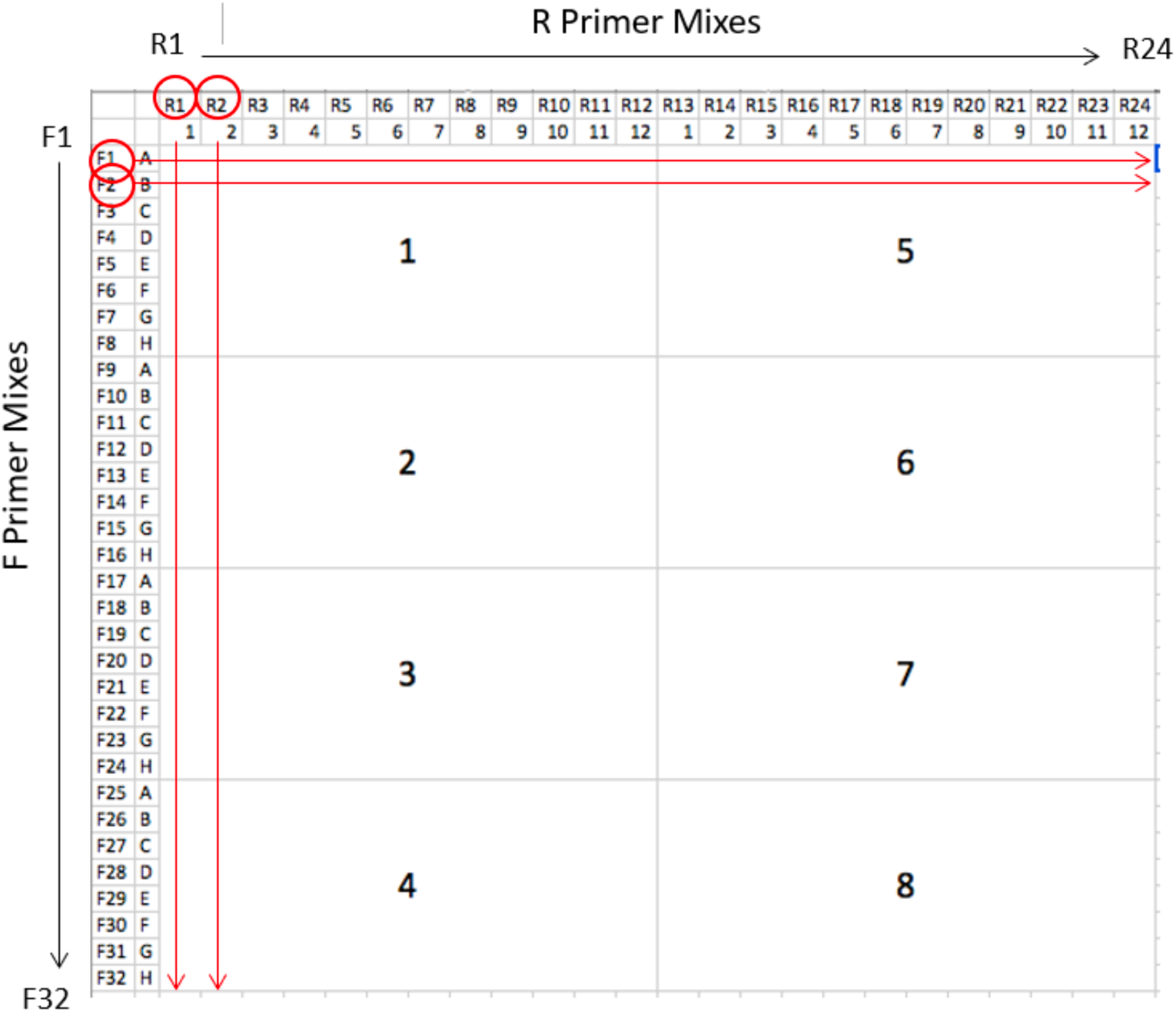

Example of preparation of 8 *Primer plates* (1-8) with 32 forward (F) primer tags and 24 reverse (R) primer tags allowing for 32 x 24 = 768 unique tag combinations (Taberlet et al., 2018). Forward and reverse primers of multiplexed markers carrying the same tags are pooled into separate 32 F primer mixes and 24 R primer mixes (2x concentrated) (i.e. one mix for each forward tag containing all forward primers and one mix for each reverse tag containing all reverse primers). These primer mixes are distributed into 96-well plates to make *Primer plates* (1x concentrated), so that each well has a unique combination of forward and reverse tagged primers for all markers in the multiplex.

### Data analysis for gray wolf, Eurasian lynx and brown bear marker panel validation

#### Methods

Genotypes of each species were organized in a custom Microsoft Access database that was used to determine consensus genotypes at each locus and perform matching of sample genotypes. Consensus genotypes at a locus for a sample were determined based on STR sequence alleles observed across PCR replicates, requiring that an allele be observed at least twice for heterozygotes and three times for homozygotes. Sample genotypes with consensus genotypes determined at ≥50% of the loci were assigned to individuals accounting for locus mismatches between similar genotypes (Paetkau 2003) and probability of identify (Waits, Luikart & Taberlet, 2001) (see (Skrbinšek et al., 2019) for details). When multiple samples were analyzed for an individual, the sample genotype with the highest QI (Miquel et al., 2006) was selected as the reference genotype for the individual. In case of missing data, the reference genotype was completed with locus-specific data from the other sample genotypes assigned to that individual.

For each species STR panel, we assessed (i) locus amplification success, as the average proportion of locus PCR yielding reads assigned to at least one allele; (ii) locus genotyping success, i.e. the proportion of consensus genotypes obtained for each locus; (iii) matching probabilities (Waits, Luikart & Taberlet, 2001); (iv) number of locus differences between individual multilocus genotypes; and (v) per locus genotyping error rates for allelic dropout and false alleles by comparing genotypes of sample PCR replicates to the individual reference genotype (Broquet & Petit, 2004).

Cumulative matching probabilities for the market panel were derived by calculating the probability of identity among siblings (PIDsibs) (Waits, Luikart & Taberlet, 2001) for each marker using program Genalex (Peakall & Smouse, 2006).

Genetic diversity was quantified by calculating the number of alleles per locus (Na), observed heterozygosity (Ho), and expected heterozygosity (He) with the *adegenet* R package (Jombart, 2008). Deviations from Hardy–Weinberg equilibrium (HWE) were tested for each locus using exact tests with 999 permutations implemented in *pegas* (Paradis, 2010). Resulting p-values were adjusted for multiple comparisons using Holm-Bonferroni sequential correction (Holm, 1979).

Pairwise linkage disequilibrium between all locus pairs was assessed in the R package *poppr* (Kamvar, Tabima & Grünwald, 2014) using the standardized index of association (*r̄_d_*). Statistical significance was evaluated with 999 permutations, and P-values were adjusted for multiple comparisons using the Holm-Bonferroni correction.

Null allele frequencies and associated confidence intervals were estimated using *PopGenReport* (Adamack & Gruber, 2014). Loci with estimated null allele frequencies greater than 0.2 were flagged as potentially affected by null alleles (Dakin & Avise, 2004).

#### Results

The wolf STR loci had an average amplification success of 68.8% across all samples. One locus (Cl434) had only 5% amplification success in the validation run and was excluded from subsequent analyses. Locus genotyping success for the other 43 loci ranged 37-100% (mean 88%) (Table 1), with 100% of the samples being genotyped at ≥50% (22) of the loci (Table S2). All samples were correctly assigned to the known individual. Individual genotypes differed at 13-41 loci and cumulative PIDsibs for the STR panel was 2.44e-15. Genotyping error rates were 16% for ADO and 1.8% for FA (Table 1).

Wolf individual genotypes had 6.5 mean alleles per locus (range 3–12), mean observed heterozygosity Ho = 0.61 and expected heterozygosity He = 0.66 (Table 1). Observed allele size range for all markers was 28-92 bp. Out of the 281 alleles identified across all loci, 57 exhibited size homoplasy within loci, sharing identical fragment lengths but differing in the underlying sequence, due to SNPs in the repeat unit or the flanking regions (Table S5). These alleles formed 28 distinct sets of same-size variants, each comprising 2–3 sequence variants, and were distributed across 17 loci, with 1–4 sets observed per locus. Five loci (Cl218, Cl251, Cl322, Cl324, Lup23) showed significant deviations from HWE and no departures from linkage equilibrium were observed after Holm-Bonferroni correction (P < 0.05; Table S6, Table S7). Observed null allele frequencies were below <0.2 except for 4 loci (Cl218, Cl251, Cl324, Lup23, Table S6).

The lynx STR loci had an average amplification success of 80% across all samples. Locus genotyping success for the 41 loci ranged 62-100% (mean 95%) (Table 1), with 100% of the samples being genotyped at ≥50% (21) of the loci (Table S3) and used for individual identification. All samples were correctly assigned to the known individual. Individual genotypes differed at 15-39 loci and cumulative PIDsibs for the STR panel was 2.4e-12. Genotyping error rates were 9.71% for ADO and 0.34% for FA (Table 1).

Lynx individual genotypes had 4.3 mean alleles per locus (range 2–9), mean observed heterozygosity Ho = 0.49 and expected heterozygosity He = 0.57 (Table 1). Observed allele size range for all markers was 41-105 bp. Out of the 177 alleles identified across all loci, 18 exhibited size homoplasy within loci, sharing identical fragment lengths but differing in the underlying sequence, due to SNPs in the repeat unit or the flanking regions. These alleles formed 9 distinct sets of same-size variants, each comprising 2 sequence variants, and were distributed across 6 loci, with 1-2 sets observed per locus (Table S5). Five loci (LL0033, LL0045, LL0087, LL0117, and LL0161) showed significant deviations from HWE, whereas no departures from linkage equilibrium were detected after Holm-Bonferroni correction (P < 0.05; Table S6, Table S7). Estimated null allele frequencies were <0.2 for all loci except LL0033, LL0045, LL0087, LL0117, and LL0206 (Table S6). When the analysis was restricted to Dinaric lynx only (n = 16), excluding translocated individuals from the Carpathian population and offsprings born after reinforcement, the patterns differed. No loci deviated significantly from HWE after Holm-Bonferroni correction (P < 0.05; Table S6) and the loci with estimated null allele frequencies exceeding 0.2 changed, with LL0045, LL0051, LL0102, LL0118, and LL0202 showing values above this threshold (LL0045 being the only locus in both datasets, Table S6).

The bear STR loci had an average amplification success of 65% across all samples. Locus genotyping success for the 30 loci ranged 15-94% (mean 76%) (Table 1), with 91% of the samples being genotyped at ≥50% (15) of the loci (Table S4) and used for individual identification. All these samples were correctly assigned to the known individual. Individual genotypes differed at 20-29 loci and cumulative PIDsibs for the STR panel was 1.04e-12. Genotyping error rates were 25.43% for ADO and 1.18% for FA (Table 1).

Bear individual genotypes had 7.7 mean alleles per locus (range 2–12), mean observed heterozygosity Ho = 0.75 and expected heterozygosity He = 0.75 (Table 1). Observed allele size range for all markers was 36-110 bp. Out of the 231 alleles identified across all loci, 66 exhibited size homoplasy within loci, sharing identical fragment lengths but differing in the underlying sequence, due to SNPs or indels in the repeat unit or the flanking regions. These alleles formed 29 distinct sets of same-size variants, each comprising 2-4 sequence variants, and were distributed across 15 loci, with 1-5 sets observed per locus (Table S5). No significant deviations from HWE and linkage equilibrium were detected after Holm-Bonferroni correction (P < 0.05) (Table S6, Table S7). Observed null allele frequencies were below <0.2 for all loci (Table S6).

